# Does the size of termite mounds affect chances of fire survival? A study of two southern African mound-building termite (Termitidae) species

**DOI:** 10.1101/2022.07.08.495810

**Authors:** B. de la Fontaine, S. Edwards

## Abstract

The effect of fire on insects is an under-researched area of ecology, especially in southern Africa. This study focusing on two termite species, *Trinervitermes trinervoides* and *Amitermes* sp., aims to determine whether fire survival in termites is related to mound size. 207 termite mounds were sampled across four sites on the periphery of a semi-urban area in Makhanda, South Africa. Each sampling site consisted of a pair of plots, either recently burnt (< 1 year or ~2 years since fire) or unburnt (> 2 years since fire). Mound type, dimensions, the presence of termites and termite species were recorded. Live mounds were significantly larger than dead mounds at both burnt and unburnt plots. For at least one termite species, there was a strong relationship between survivorship and size class of mounds at burnt plots but not at unburnt plots, suggesting that mound size plays a role in fire survival.

## INTRODUCTION

Mound-building termites (Blattaria: Termitidae) play a beneficial role in savannas, grasslands, and tropical forests as invertebrate decomposers of plant matter (Trapnell *et al*. 1976; Davies *et al*. 2010) and as landscape architects or ecosystem engineers (Dangerfield *et al*. 1998; Hölldobler and Wilson, 2009). In savannas in particular, termites contribute to nutrient cycling and soil water retention in a habitat where both soil nutrients and water are limiting factors for plant growth to a greater extent than most other habitat types (Sage and Kubien, 2003; Davies *et al*. 2009). Ants (Hymenoptera: Formicidae), lacking the cellulose-digesting, endosymbiotic microorganisms that have co-evolved with termites, are not physiologically capable of providing the same ecosystem service to the same extent, although some South American ants and African termites utilize ectosymbiotic, cellulose-digesting fungi (Aanen *et al*. 2002). However, both termites and ants contribute to soil turnover and water retention through tunnelling behaviour in such a way that has been shown to increase crop yields in arid climates by a large margin (Evans *et al*. 2011).

It is in the interests of ecologists and landowners in fire-prone regions to understand the positive and negative impacts of fire on termites. Despite the ecological importance of invertebrates in ecosystems that naturally experience frequent fire regimes, namely savanna-grasslands, few studies to date have attempted to examine the fire-invertebrate dynamic in these ecosystems (Davies *et al*. 2010), and even fewer studies have been carried out in southern Africa (Uys *et al*. 2006; Davies *et al*. 2012). The diversity of this region is often overlooked: for instance, *Trinervitermes trinervoides*, a species distributed across southern Africa and common in South Africa, is curiously absent from catalogues such as Ferra?s (1982) series *Termites of a South African Savanna* and Krishna and coworkers’ treatment of Termitidae in their *Treatise on the Isoptera of the World* (Krishna *et al*. 2013).

Termites have been suggested to be resistant to fire to varying degrees based on studies conducted in Australian (Abensperg-Traun *et al*. 1996; Dawes-Gromadzki *et al*. 2007; Avitabile *et al*. 2015), West African (Benzie, 1986), Brazilian (DeSouza *et al*., 2003) and South African (Davies *et al*. 2012) fire biomes. These studies invariably examine the effects of fire in terms of termite abundance and/or diversity, but do not provide a great amount of information on the fate of individual colonies, which will inevitably be a difficult task with non-mound-building species. More importantly, these studies do not take into consideration or attempt to test for a possible interaction between mound size and fire. In *T. trinervoides*, mound size is negatively correlated with monthly temperature variability; alates and brood are kept at the core of the mound where temperatures for growth are ideal (Field and Duncan, 2013). Mound size, then, is expected to influence the degree to which termites are able to withstand thermal extreme. A study by Davies *et al*. (2012) in the Kruger National Park represents a distinct ecosystem from the Eastern Cape grasslands and does not account for the two dominant mound-building species in this ecoregion, whose different styles of mound construction may affect their responses to fire.

In April and July of 2021, fire burned areas of grassland and veld in Makhanda (formerly Grahamstown), South Africa, affecting several thousand termite mounds. Two mound-building termite species occur in this area: *Trinervitermes trinervoides*, with nasute soldiers, and an *Amitermes* species (possibly *Amitermes hastatus* or a closely related species; any use of *“Amitermes* sp.” hereafter refers to this unidentified species), whose soldiers have pincer-like jaws and are much less numerous in the mound than those of the former. The two species construct mounds that are distinctly different in shape, size and hardness and often in colouration (even on the same soil), using slightly different materials (Fig. 1). *Trinervitermes trinervoides* mounds are consistently taller, rounder, darker in colour, and constructed of a more brittle combination of clay and grass litter, which is stocked in small chambers in the outermost layers of the mound and also incorporated into the structure of the mound exterior. Grass litter is the primary food source of *T. trinervoides*, earning them the name “harvester termites”. The species is considered to be the primary invertebrate decomposer in southern Africa’s semi-arid grasslands (Adam *et al*. 2018). *Amitermes* sp. mounds are flatter and more irregular in shape and are in most cases constructed of very fine clay or silt, making them very solid structures. *Amitermes* sp. are soil- and wood-feeders and do not typically store or incorporate harvested plant material in their mound structures, although they often construct their mounds around tree stumps, and infrequently will construct small, dark, and brittle carton-mounds composed largely of woody material from decaying stumps (personal observation). The two termite species often construct their mounds directly adjacent to each other (as close as 10 cm – Fig. 1), much closer than is normally tolerated for conspecific mounds of other colonies (Theron, 2013). Low interspecific aggression is to be expected, since the two species are unlikely to compete for the same food resources. Subterranean termites (Hodotermitidae) are also common in this area but were not sampled.

**Figure 1.**
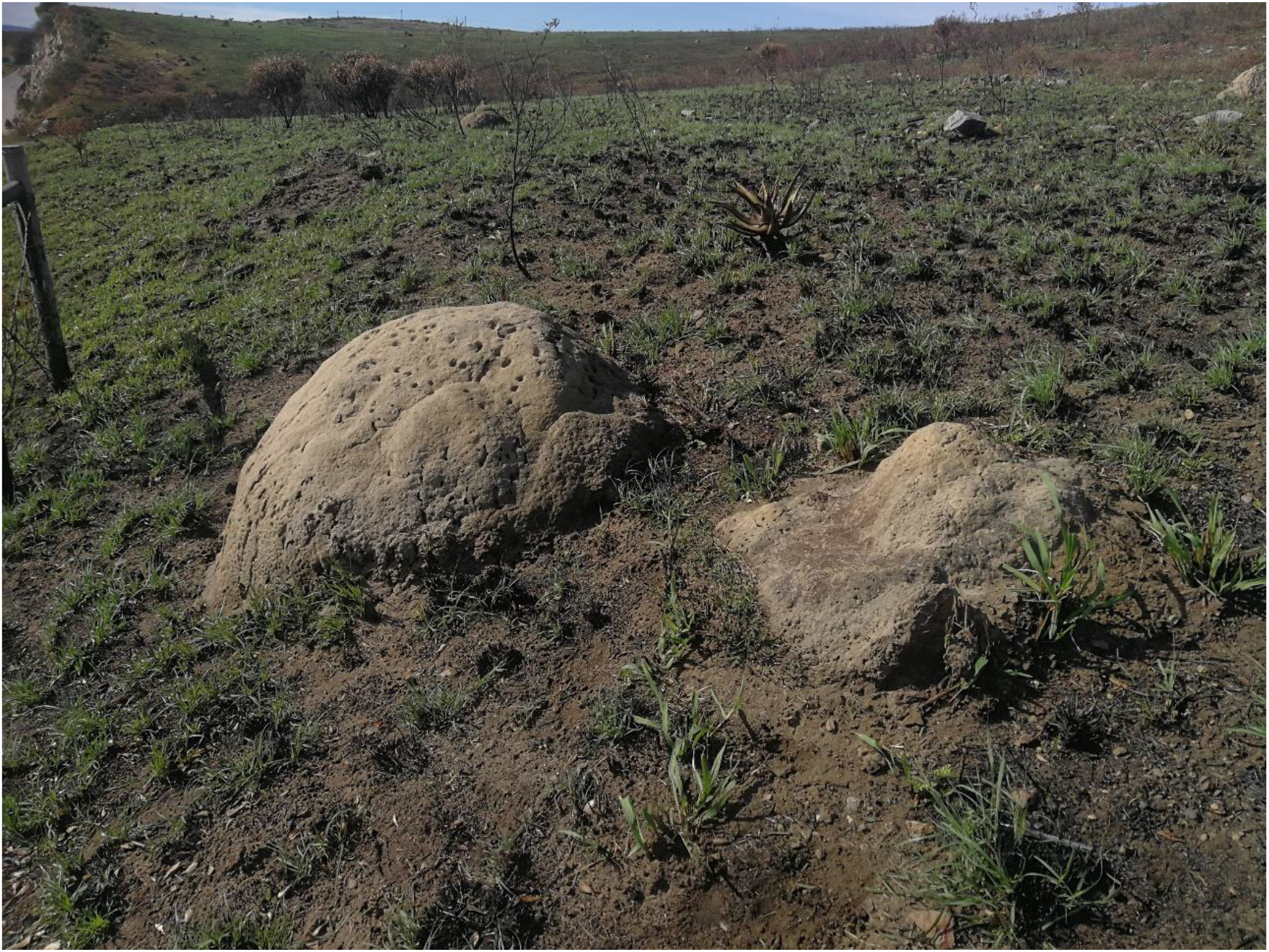
Mounds of the termite species *T.trinervoides* (left) and *Amitermes* sp. (right) on recently-burnt grassland. The two species have noticeably different styles of mound construction, with mounds of the *T. trinervoides* being distinctly taller, more regularly dome-shaped, more brittle and often darker. The two species may construct their mounds within as little as half a metre of each other.

Here we investigate whether or not there is a relationship between mound size and fire survival in *T. trinervoides* and *Amitermes* sp. by surveying mounds at different time intervals after fire.

## MATERIALS AND METHODS

A total of 207 termite mounds were sampled from July to August 2021 at four sites (Sites A, B, C and D) on the periphery of semi-urban Makhanda. Each site consisted of one burnt plot (< 1 year or ~ 2 years after fire) and an adjacent unburnt plot (~> 2 years after fire) (Table 1). Burnt plots were identified by personal observation of active fires or statements from residents, and were distinguished from unburnt plots by the presence of burnt vegetation, rock varnishes, and differences in the average height of bunch grasses and woody plants. Recently burnt plots (< 1 year after fire) were sampled within the four-month post-burn period during which the effects of fire on vegetation remain significant, according to Davies *et al*. (2012).

**Table 1.**
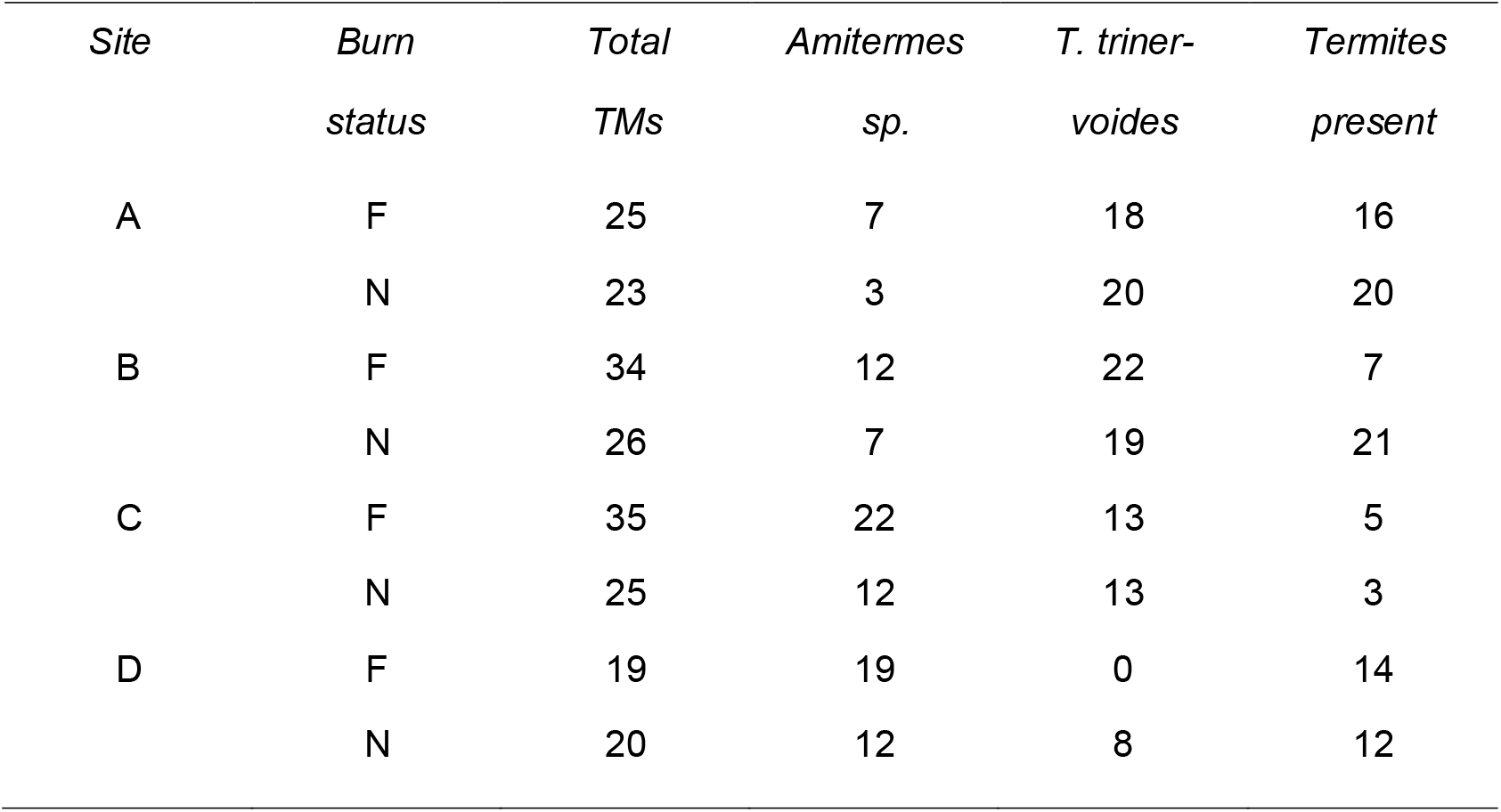
Tallies of termite mounds (TMs) from sites A-D, divided into burnt (F) and unburnt (N) plots. Species names indicate the species that constructed the mound, regardless of whether termites were present.

| *Site* | *Burn status* | *Total TMs* | *Amitermes sp.* | *T. triner-voides* | *Termites present* |
| --- | --- | --- | --- | --- | --- |
| A | F | 25 | 7 | 18 | 16 |
|  | N | 23 | 3 | 20 | 20 |
| B | F | 34 | 12 | 22 | 7 |
|  | N | 26 | 7 | 19 | 21 |
| C | F | 35 | 22 | 13 | 5 |
|  | N | 25 | 12 | 13 | 3 |
| D | F | 19 | 19 | 0 | 14 |
|  | N | 20 | 12 | 8 | 12 |

Sites A and B include plots which burned in April and July of 2021, respectively. Site A (Mountain Drive – 33°19’57”S 26°31’47”E) is part of a large indigenous grassland formerly invaded by alien trees (*Pinus* and *Acacia* spp.) and since restored through a community tree-felling initiative. Many logs and stumps still remain, which together with the abundance of grass litter contributes to the very high density of both termite species. The area is often grazed by free-roaming cattle. Site B (Settler’s Monument – 33°19’04”S 26°31’04”E), located near a national monument and convention centre, consists of mixed grassland and low-density thicket, with some planted trees that are not native to the area.

Sites C and D include plots which burned in 2019. Site C (Cradock Heights – 33°17’53.4”S 26°30’21.5”E) is located near a residential area and ranges from scrub in the burnt plot to dense thicket in the unburnt plot. Site D (Makhanda Aerodrome – 33°17’24.6”S 26°30’23.5”E) is located near an airfield and consists of mixed grassland and scrub on rocky ground.

The following information was recorded for each termite mound: maximum diameter and height in cm; presence or absence of termites (mound “live” or “dead”); the termite species that constructed the mound; and the structural state of the mound. Mound dimensions were measured to one decimal place with a graduated tape measure. The spherical cap equation was used to estimate the volume of mounds, assuming all mounds to be roughly dome-shaped:

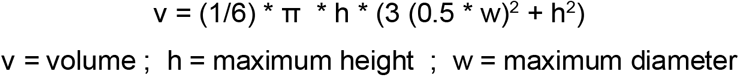

The presence or absence of termites was determined by making one to three small holes (~6 cm diameter) in the exterior of the mound and by examining the structural state of the mounds (i.e. in good shape or clearly degraded).

Data were analysed in R Studio 1.4.1 (R v.4.0.4) (R Core Team, 2021). Pearson’s Chi-squared test (R Base, function *chisq.test)* was used to compare differences in the numbers of live versus dead mounds by species, site burn status and time-since-fire. Differences in mound size by species and site burn status were compared using Wilcoxon Rank Sum tests (R Base, function *wilcox.test)*.

In order to disentangle the relationship between fire and mound size from other influencing factors, mounds of each species were grouped into five size classes (ranging from very small to very large) (Table 2). Linear regression (R Base, function *lm*) was applied to determine the relationship between mound size class and the proportion of live mounds which were from burnt plots (“fire survivors”).

**Table 2.** Explanation of mound size classes for *Amitermes* sp. and *T. trinervoides*. # indicates number of mounds per size class for each species. Sizes are in cm^3^.

| *Size classes* | *Amitermes* | *#* | *T. trinervoides* | *#* |
| --- | --- | --- | --- | --- |
| Very small | <1000 | 4 | <5000 | 10 |
| Small | <20,000 | 24 | <15,000 | 23 |
| Medium | 20,000-50,000 | 38 | 15,000-55,000 | 57 |
| Large | >50,000 | 23 | >55,000 | 23 |
| Very large | >120,000 | 5 | >200,000 | 4 |

## RESULTS

At sites sampled < 1 year after fire (Sites A and B), unburnt plots had a significantly greater proportion of live mounds versus dead mounds than burnt plots, but there was not a significant difference at sites sampled ~ 2 years after fire (Sites C and D) (Table 2). The proportion of mounds by either species differed between sites with different time-since-fire: mounds constructed by *T. trinervoides* made up the greater proportion of mounds at Sites A and B (40 of 59) (χ2 = 7.475, df = 1, p = 0.0063) while mounds of *Amitermes* sp. were predominant at Sites C and D (41 of 54) (χ2 = 14.519, df = 1, p < 0.001).

Live mounds were significantly larger than dead mounds at burnt plots but not at unburnt plots for both species. Live mounds at burnt plots were on average almost three times larger on average in *Amitermes* sp. and more than twice as large in *T. trinervoides* (Table 3).

**Table 3.** Results of statistical tests on count data at burnt and unburnt plots. Statistically significant p-values are indicated in bold.

| *Time-since-fire* | *Chi-squared test* | *No. of live mounds* |
| --- | --- | --- |
|  | *(burnt vs. unburnt plot)* |  |
| < 1 yr | $\chi^2 =$ | 5.0625 |
|  | df = | 1 |
|  | p = | **0.03** |
| ~ 2 yrs | $\chi^2 =$ | 0.471 |
|  | df = | 1 |
|  | p = | 0.49 |

**Table 4.**
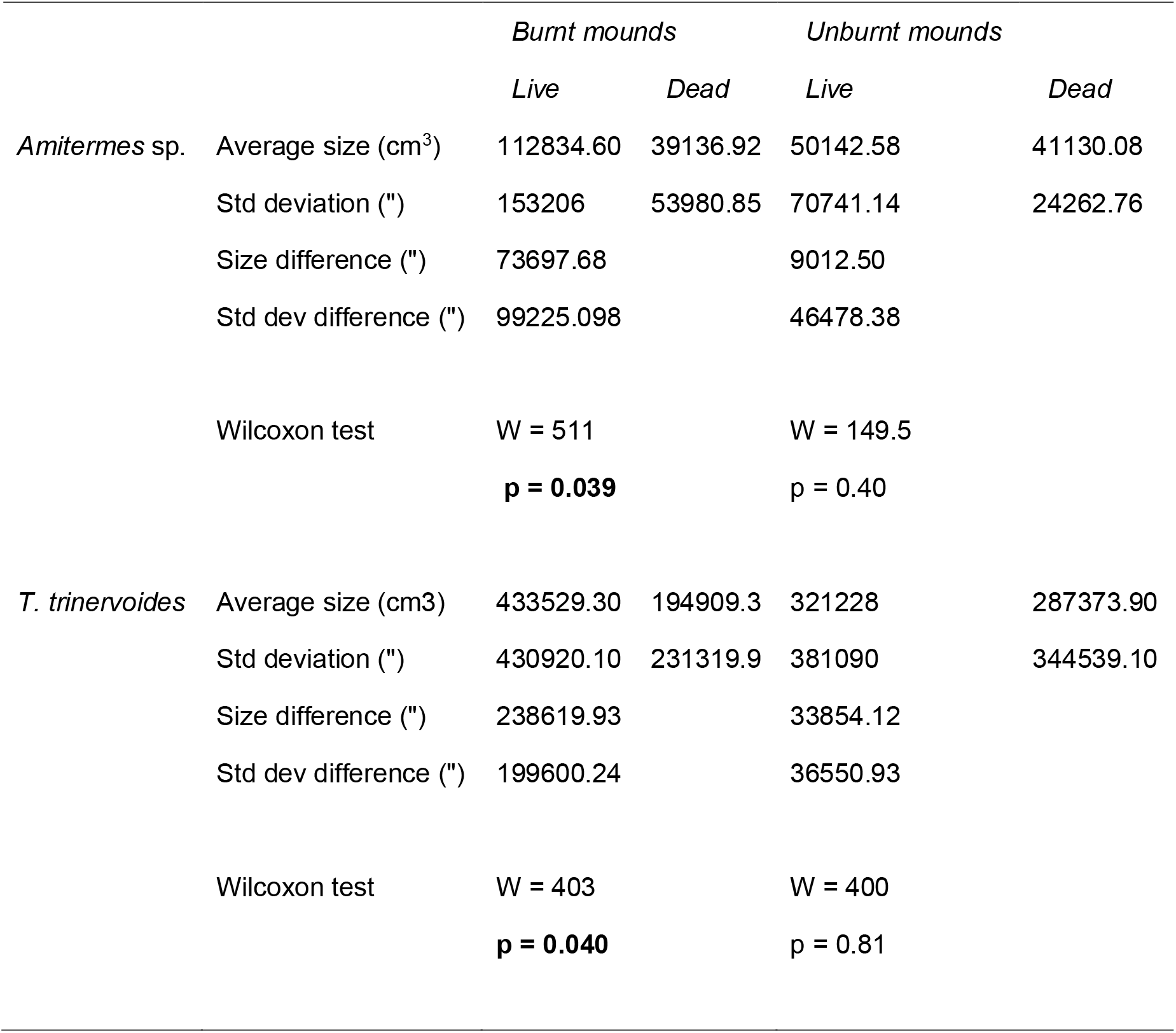
Results of statistical tests on mound size measurements at burnt and unburnt plots. Statistically significant p-values are indicated in bold.

There was a positive relationship between size class and fire survivorship for mounds of *T. trinervoides* (R^2^ = 0.80, F = 12.07, p = 0.040) and *Amitermes* sp. (R^2^ = 0.75, F = 8.959, p = 0.058). The pattern of the survival-size relationship was very similar in both species: a general upward trend with increasing size class, but with a dip in the medium size class (Fig. 2). The relationship between size and survivorship for mounds at unburnt plots was somewhat weaker for *T. trinervoides* (R^2^ = 0.70, F = 6.94, p = 0.078) and much weaker for *Amitermes* sp. (R^2^ = 0.25, F = 1.017, p = 0.388).

**Figure 2.**
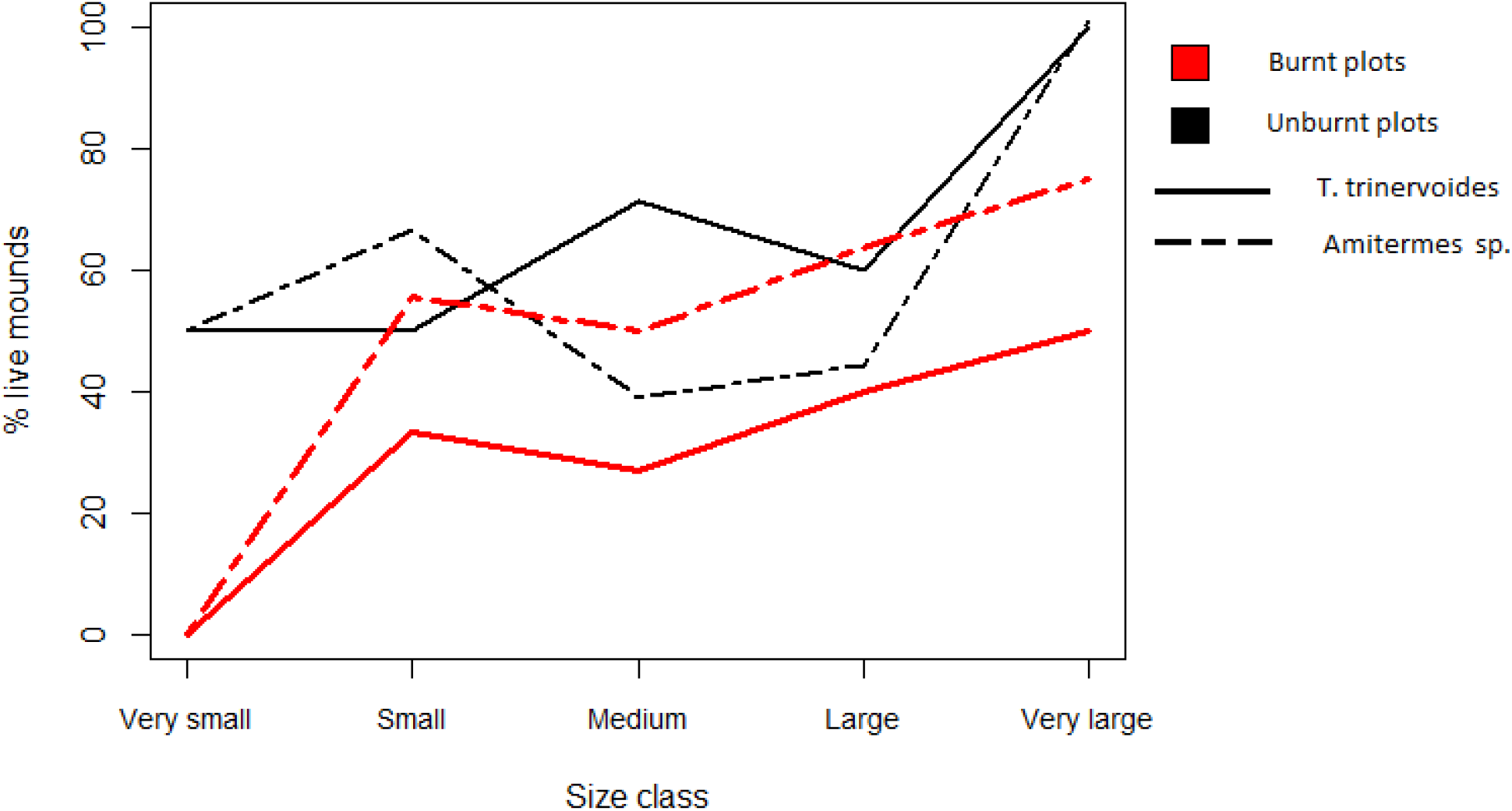
Graph showing the percentage of live mounds of each size class from burnt and unburnt plots for two termite species.

## DISCUSSION

There is a relationship between size and survival in termite mounds in which fire is a major factor. The percentage of mounds surviving after fire was strongly linked to size class in *T. trinervoides* and moderately in *Amitermes* sp., and in both termite species live mounds were, on average, larger than dead mounds, particularly at burnt plots. However, in unburnt plots, the size-survival relationship was still relatively strong in *T. trinervoides* while in *Amitermes* sp. it was effectively non-existent. This suggests that fire affects the two species differently and is more important for *Amitermes* sp. than for *T. trinervoides*. Why might this be?

Without having compared mounds pre- and post-burn, it is not clear whether differences in mound size are a consequence of thermoregulatory ability contributing to survival of extreme high temperatures (the direct effects of fire), or of larger mounds being more resilient during the food scarcity that follows fires (indirect effects). The greater proportion of dead mounds compared to live mounds in the first few months after fire indicate that fire does “kill” some termite mounds outright, although termites appeared to recover within two years. (It is not clear whether termites achieve this recovery solely by building new mounds; some researchers have suggested that termite colonies migrate during and after fires, in which case they would be capable of vacating and then recolonizing old mounds (Abensperg-Traun *et al*. 1996; DeSouza *et al*. 2003)) If the direct effects of fire are more important, differences in mound structure and functional characteristics (e.g. ventilation and insulation) may explain the difference in response between species (Abensperg-Traun & Milewski, 1995). The most obvious difference between the two species, however, is their use of food resources. *Amitermes* sp., being preferential wood-feeders, are expected to be the worst-affected by fire, especially in grassland habitats where woody vegetation is already scarce. Therefore, the longer-term decline and increase in live termite colonies mainly reflects the loss and recovery of above-ground vegetation. Said another way, the effects of fire on vegetation are more important than fire itself as a factor in termite mound survival, as concluded in several studies (Abensperg-Traun *et al*. 1996; Davies *et al*. 2012; Avitabile *et al*. 2015).

In terms of vegetation, there were some notable differences between sites and between plots within sites. Sites A and B (burned in 2021) consist of open grassland and a mix of grassland and low-density thicket respectively, while Sites C and D (burned in 2019) were more diverse, ranging from degraded scrubland with little or no grass cover to dense thicket. Thus, plots with the greatest time-since-fire had a higher abundance of woody vegetation as opposed to grasses, which affects both invertebrate composition (including termites and ants) and fire dynamics. The difference in vegetation between sites is also the most obvious reason for the differences in termite abundance and species composition: *T. trinervoides* mounds were absent from the burnt plot at Site D (too little grass cover) and the number of live termite mounds was particularly low at Site C (very little vegetation at either plot).

The possibility that larger termite mounds are more likely to survive fire and the temporary denuding of the landscape hints at a hypothetical role as “mother colonies”. These would be very large and old termite colonies that, because they are well-established and more resistant to recurring threats such as fires, seed the landscape with younger offshoot colonies after these and other extreme events kill off a large portion of the population. In this way, mother colonies could help their own species to persist in an area under harsh conditions. Damaging or removing only a few large termite mounds, either because they obstruct the path of farm equipment or because termites are perceived to be pests, might have serious long-term impacts on termite populations and ecosystem function in fire biomes.

Fire and its effects on vegetation cover are not likely to be the only factors selecting for larger mounds. A range of other environmental and biotic factors are expected to have a large influence on the chances of colony survival. These include annual rainfall, rainfall and temperature variability, predation, density-dependent competition with conspecifics and perhaps also interspecific competition. The region had also experienced drought for five consecutive years at the time the study was conducted, a major stressor regardless of any effects of fire. With the drought recently alleviated there is the potential for a fresh study to be conducted that resolves some of the issues mentioned by selecting sites that are uniform, for instance all open grassland, and monitoring the same termite mounds before and after fire over a longer period.

## ETHICS

This study was done with the permission of the Rhodes University Animal Research Ethics Commission (RUAR-EC Project ID 5109).

## DATA AVAILABILITY

The original data collected in this study are available from the corresponding author on request.

## AUTHORS CONTRIBUTIONS

SE conceived of the idea. BD and SE designed the study. BD carried out the fieldwork and data analysis. Both authors contributed to the draft manuscript and gave approval for publication.

## ACKNOWLEDGEMENTS

We thank Professor Martin Villet of Rhodes University for identifying burnt areas in Makhanda from 2019, advice about statistical analyses, and proofreading draft versions of this paper.

